# Sec22b regulates inflammatory responses by controlling the nuclear translocation of NF-κB

**DOI:** 10.1101/2020.09.20.305383

**Authors:** Guillermo Arango Duque, Renaud Dion, Aymeric Fabié, Julien Descoteaux, Simona Stäger, Albert Descoteaux

**Affiliations:** INRS – Centre Armand-Frappier Santé Biotechnologie, Université du Québec, Laval, QC, H7V 1B7, Canada

**Keywords:** Cytokine, intracellular trafficking, nuclear translocation, soluble NSF attachment protein receptor (SNARE), NF-kappa B (NF-KB)

## Abstract

Soluble NSF attachment receptor (SNARE) proteins regulate the vesicle transport machinery in phagocytic cells. Within the secretory pathway, Sec22b is an ER-Golgi intermediate compartment (ERGIC)-resident SNARE that controls phagosome maturation and function in macrophages and dendritic cells. The secretory pathway controls the release of cytokines and may also impact the secretion of nitric oxide (NO), which is synthesized by the Golgi-active inducible nitric oxide synthase (iNOS). Whether ERGIC SNARE Sec22b controls NO and cytokine secretion, is unknown. Using bone marrow-derived dendritic cells (BMDC), we demonstrated that iNOS colocalizes with ERGIC/Golgi markers, notably Sec22b and its partner syntaxin-5 (Stx5), in the cytoplasm and at the phagosome. Pharmacological blockade of the secretory pathway hindered NO and cytokine release, and inhibited NF-κB translocation to the nucleus. Importantly, RNAi-mediated silencing of Sec22b revealed that NO and cytokine production were abrogated at the protein and mRNA levels. This correlated with deregulated mitogen-activated protein kinase signalling and reduced nuclear translocation of NF-κB. We also found that Sec22b co-occurs with NF-κB in both the cytoplasm and nucleus, pointing to a role for this SNARE in the shuttling of NF-κB. Collectively, our data unveiled a novel function for the ER-Golgi, and its resident SNARE Sec22b, in the production and release of inflammatory mediators.

## Introduction

The innate immune function of professional phagocytes is channeled through their capacity to ingest and degrade foreign particles and microbes (1). Following particle binding and uptake, phagocytes release a series of small molecules such as nitric oxide (NO) and cytokines that dictate the nature and intensity of the immune response. Particularly in dendritic cells, the phagosome also serves as a platform for antigen processing and crosspresentation (2). The aforementioned processes exert a membrane demand that is actively supplied by endosomes and the secretory pathway, a dynamic group of organelles through which proteins and lipids are synthesized, processed and transported (3,4). While recycling/early endosomes and the endoplasmic reticulum (ER) supply membrane and protein effectors to nascent phagosomes (5,6), the latter also serve as cytokine secretion sites (7). Through sequential membrane exchanges with organelles such as the lysosome, maturing phagosomes acidify and become highly degradative.

Interorganellar vesicular transport is regulated by proteins that reside and determine cargo selectivity and vesicle destination (3). Vesicle docking and fusion rely on organelle-specific members of the soluble *N*-ethylmaleimide-sensitive factor attachment protein receptor (SNARE) family (4,8). For instance, the syntaxin (Stx) 4/SNAP23/VAMP3 complex facilitates the passage of TNF from recycling endosomes to the cell membrane (7,9). On the other hand, trafficking within the ER, ER-Golgi intermediate compartment (ERGIC), Golgi apparatus and phagosome circuitry is orchestrated by SNARE complexes involving Sec22b, Stx5 and Stx4 (10,11). Notably, their absence hinders phagosome maturation and antigen crosspresentation (10,11), which translates into phenotypes such as impaired bacterial and tumor clearance (12–14). Moreover, a panmouse knockout of Sec22b is embryonically lethal (15), highlighting the key importance of this ER/ERGIC-resident SNARE in immunity and development.

The inducible nitric oxide synthase 2 (iNOS) bridges homeostatic regulation and immunity via the production of NO from L-arginine (16). NO modulates inflammation, vasodilation and neurotransmission. Importantly, this reactive molecule contributes to the generation of highly microbicidal reactive nitrogen species (RNS) within the phagosome (17,18) that kill internalized bacteria and *Leishmania* parasites (19). iNOS expression is mainly regulated at the transcriptional level by nuclear factor kappa B (NF-κB), and enzymatic activity is regulated at the translational and post-translational levels. Indeed, iNOS has to assemble into homodimers in order to transform L-arginine into RNS (20). Although its intracellular trafficking remains uncharacterized, iNOS-positive vesicles have been observed to reside in ER-associated aggresomes and the trans-Golgi (21), as well as the phagosome surface (17). Therefore, the question emerges as to whether secretory pathway-resident SNAREs regulate iNOS trafficking and function. We showed that iNOS is mainly localized to the ERGIC/Golgi both in the cytoplasm and at the phagosome. Notably, Sec22b knockdown hindered NF-κB nuclear translocation, thereby decreasing iNOS and cytokine production. We also observed that Sec22b and NF-κB co-occur in the cytoplasm and the nucleus. These findings highlight a new role for this SNARE in the control of inflammatory responses.

## Results

### iNOS localizes within the secretory pathway in BMDC and is recruited to the phagosome

In BMDC, the release of NO is induced by both soluble and particulate stimuli (Figure 1A). To elucidate the underlying mechanisms and pathways, we first assessed the localization of iNOS in LPS-stimulated BMDC, using immunofluorescence confocal microscopy. Figure 1B shows that iNOS was present in vesicular structures situated in the cytoplasm and plasmalemma, consistent with its role in NO secretion to the extracellular milieu (16). To further determine the subcellular localization of iNOS in BMDC, we performed co-localization analyses by immunofluorescence confocal microscopy using markers of the ER (CNX, PDI), the ERGIC (ERGIC53, Sec22b, Stx5) the Golgi apparatus (P115) and lipid rafts (CTX) (Figure 1C, D and Figure S1). These analyses revealed that iNOS was widely present in secretory pathway organelles, though most saliently in the ERGIC and the Golgi (Figure 1C, D). Moreover, following phagocytosis of zymosan particles, we observed that iNOS colocalizes with Sec22b, Stx5, and LAMP1 on phagosomes (Figure 2A, B). Collectively, these data suggest a role for the secretory pathway in NO secretion.

**Figure 1.**
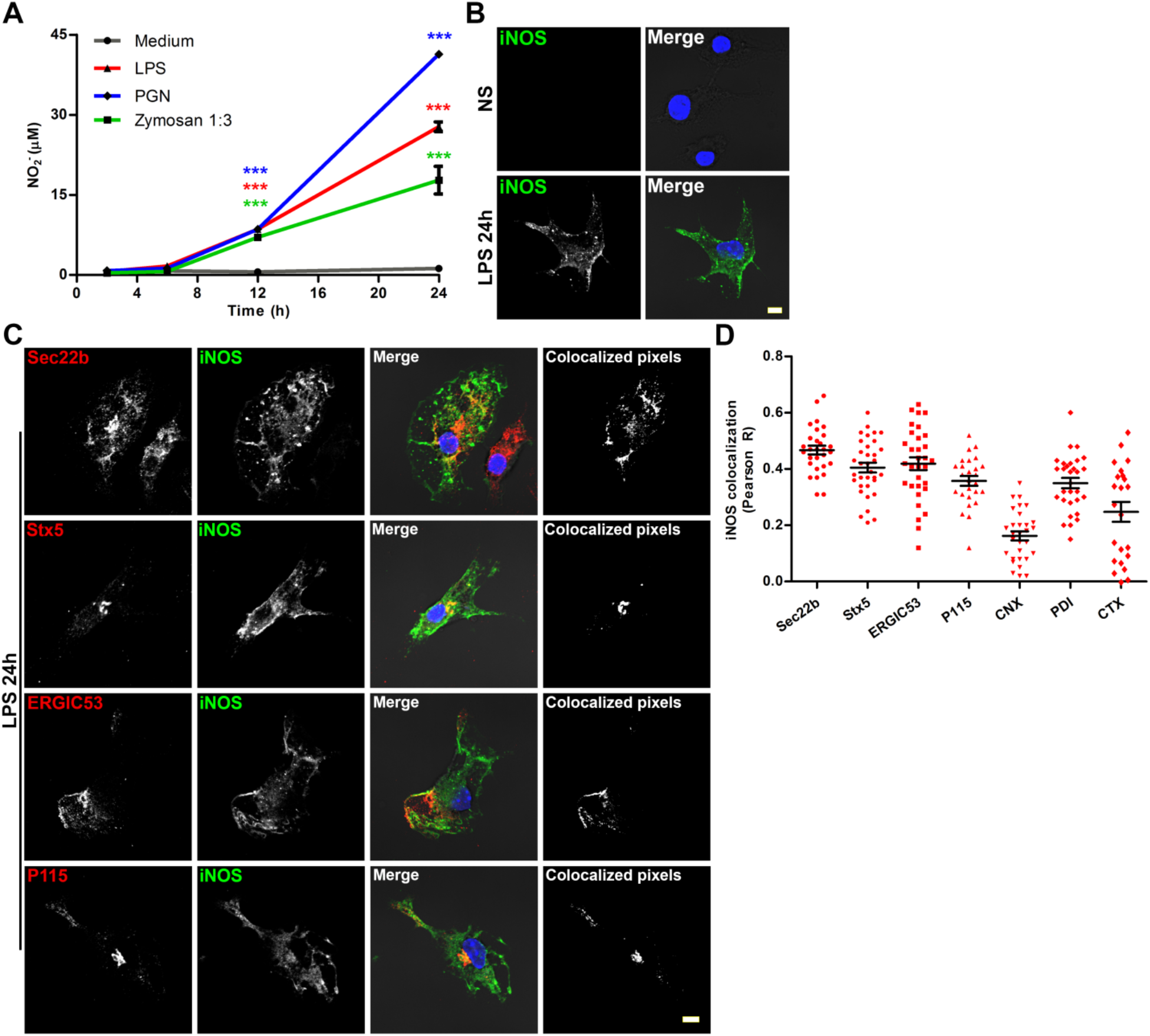
iNOS colocalizes with ERGIC and Golgi markers. **(A)** BMDC were incubated with LPS, PGN and zymosan and NO release was assessed by measuring NO_2_^-^ over a period of 2-24 h. Data are presented as mean concentration ± SEM of three independent experiments done in triplicate wells. ***, *p* < 0.001. **(B)** The intracellular expression of iNOS (green) in non-stimulated (NS) or LPS-stimulated BMDC was visualized via confocal immunofluorescence microscopy. DNA is in blue; bar, 5 μm. **(C)** BMDC were stimulated with LPS for 24 h and the colocalization (white pixels, rightmost panels) of iNOS (green) with ER/ERGIC-associated proteins Sec22b, Stx5 (red); ERGIC maker ERGIC53 (red); and Golgi marker P115 (red) was assessed by immunofluorescence. DNA is in blue; bar, 5 μm. Images are representative of three independent experiments. **(D)** iNOS colocalization was quantified using the Pearson method (see also Figure S1). Data are presented as mean Pearson R coefficient ± SEM of three independent experiments (≥10 cells per experiment); each point represents the coefficient of a single cell.

**Figure 2.**
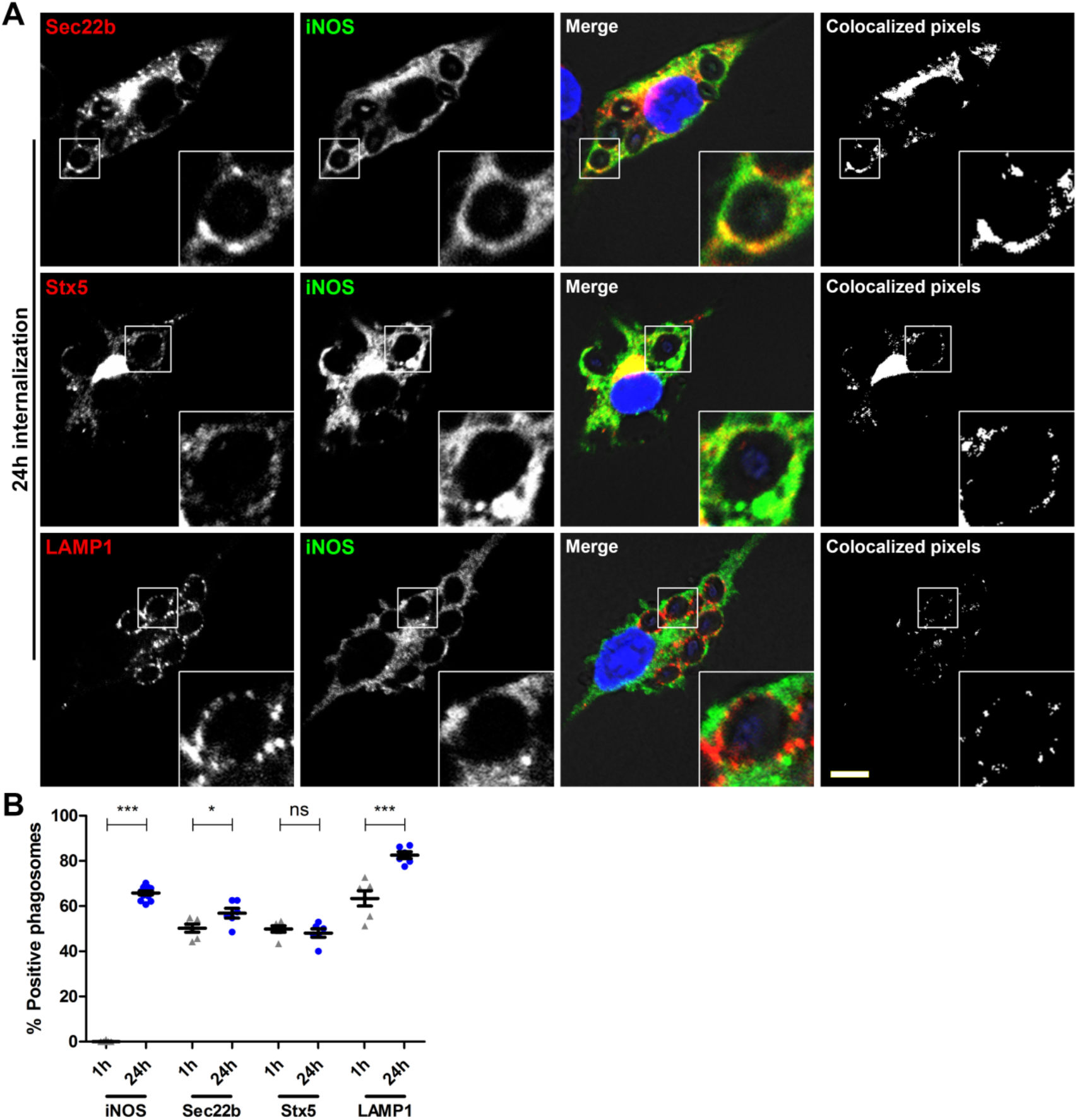
iNOS, Sec22b and Stx5 are recruited to the phagosome. **(A)** BMDC were fed with opsonised zymosan and recruitment of iNOS (green), Sec22b (red) and Stx5 (red) was visualized via immunofluorescence 24 h post-internalization. Recruitment of LAMP-1 (red) was used as reference. Cytoplasmic and phagosomal (3X-enlarged insets) colocalization can be observed in the rightmost panels (white pixels). DNA is in blue; bar, 5 μm. Images are representative of three independent experiments. **(B)** Phagosomal recruitment quantification of the aforementioned proteins at 1 h and 24 h post-phagocytosis. Data are presented as mean % positive phagosomes ± SEM of three independent experiments done in triplicate coverslips; each point represents % recruitment in 100 phagosomes per coverslip. *, *p* < 0.05; ***, *p* < 0.001; ns, not significant.

### The secretory pathway is required for NO and cytokine production

The ER and the Golgi are critical for the biogenesis and secretion of proinflammatory cytokines such as TNF and IL-6 (4). Here, we assessed the potential role of ER-Golgi trafficking in the control of iNOS activity and NO release by BMDC, and we included TNF and IL-6 in our analysis since their release is mediated by the secretory pathway (4). To this end, we treated BMDC with DMSO or with inhibitors of ER-Golgi trafficking, namely brefeldin A, monensin, and retro-2 (22–24), prior to stimulation with LPS. While monensin and brefeldin A inhibited NO, TNF and IL-6 release, retro-2 only inhibited TNF release, highlighting individual differences in the intracellular trafficking of those molecules (Figure 3A-C). Of note, both monensin and brefeldin A inhibited iNOS protein expression (Figure 3D-E), suggesting that ER-Golgi trafficking is part of the pathway leading to iNOS expression. To understand how perturbation of the secretory pathway led to the inhibition of NO and cytokine release, we first assessed the impact of monensin on the integrity of LPS-induced signal cascades associated to the expression of iNOS, TNF, and IL-6 (25,26). As shown in Figure 4A, pretreatment with monensin had no effect on LPS-induced phosphorylation of p44/p42 MAPK (ERK1/2) and degradation of IκBα, but impaired the phosphorylation kinetics of p38 and JNK. One of the key downstream event of inducible IκBα degradation is the nuclear translocation of the transcription factor NF-κB, which mediates the expression of a large number of LPS-inducible genes including iNOS, TNF, and IL-6 (25,26). As shown in Figure 4B-C, pretreatment of BMDC with monensin impaired LPS-induced nuclear translocation of NF-κB. Collectively, these findings are consistent with the notion that the integrity of the secretory pathway is necessary for optimal NF-κB-mediated LPS responses in BMDC.

**Figure 3.**
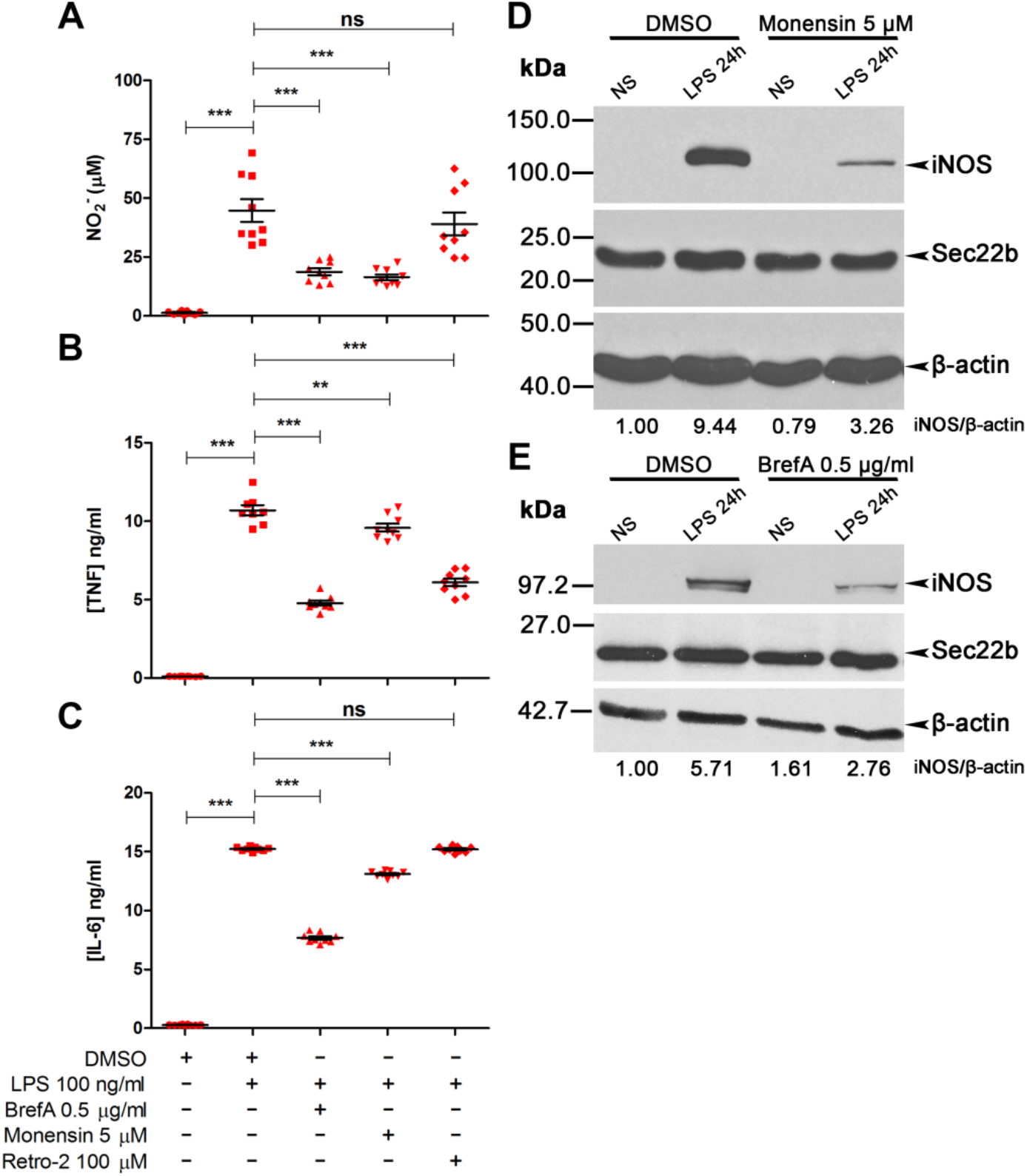
Inhibition of ER-Golgi trafficking blocks the production of inflammatory mediators. BMDC were incubated with DMSO or the indicated pharmacological inhibitors for 2 h prior to LPS stimulation for 24 h. Cell culture supernatants were then assayed for NO **(A)**, TNF **(B)** and IL-6 **(C)** release. Data are presented as mean concentration ± SEM of three independent experiments done in triplicate wells; each point represents the concentration of a well. **, *p* < 0.01; ***, *p* < 0.001; ns, not significant. Immunoblotting was used to assay iNOS levels in non-stimulated (NS) or LPS-stimulated cells treated with brefeldin A (BrefA) **(D)** or monensin **(E)**. iNOS densitometries were normalized with those of the β-actin loading control, and expressed relative to non-stimulated DMSO-treated cells. Images are representative of at least two experiments.

**Figure 4.**
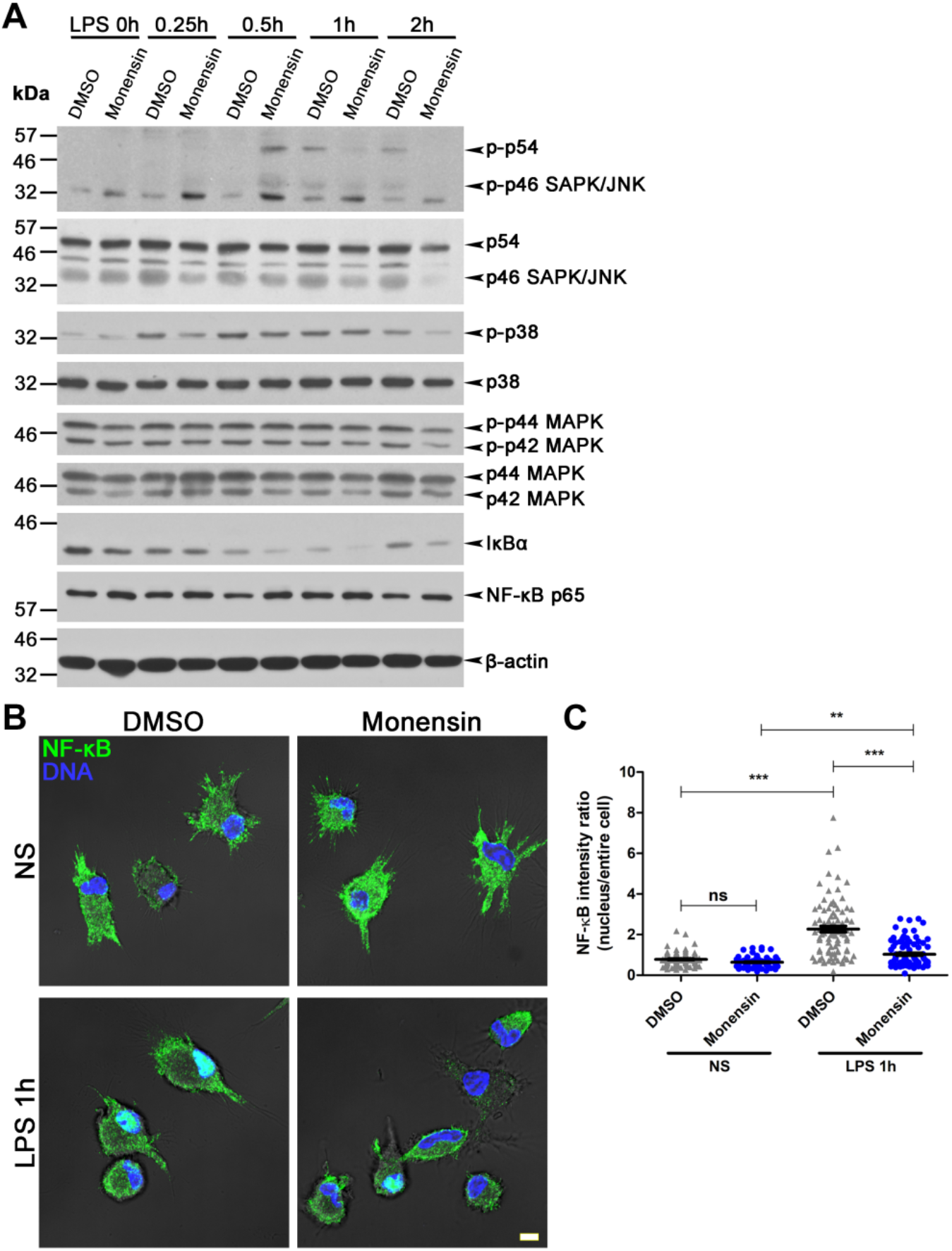
Inhibition of ER-Golgi trafficking inhibits MAPK phosphorylation and the translocation of NF-κB to the nucleus. **(A)** BMDC were incubated with DMSO or monensin for 2 h prior to LPS stimulation for 0.25-2 h. Western blots show levels of total and phosphorylated MAPK, as well as IκBα and NF-κB. β-actin was used as loading control and images are representative of two experiments. **(B)** DMSO- or monensin-treated BMDC were incubated with LPS for 2 h, and nuclear translocation of NF-κB (green) was visualized and quantified **(C)** via immunofluorescence. DNA is in blue; NS, non-stimulated; bar, 5 μm. Data are presented as average MFI ratios (NF-κB MFI of nucleus/entire cell) ± SEM of four independent experiments (≥20 cells per experiment); each point represents the ratio of a single cell. **, *p* < 0.01; ***, *p* < 0.001; ns, not significant.

### The secretory pathway SNARE Sec22b regulates LPS-induced responses

The SNARE Sec22b plays a role in ER-Golgi trafficking (27) and in the unconventional secretion of IL-1β in cells treated with autophagy inducers, a process known as secretory autophagy (28). To assess the potential role of Sec22b in the release of NO, we stimulated with LPS DC-like JAWS-II cells transduced with an shRNA to Sec22b (shSec22) or a scrambled (shScr) sequence (10). As shown in Figure 5A-C, Sec22b knockdown diminished the release of NO, as well as of TNF and IL-6. To confirm the role of Sec22b in NO secretion and to test whether this mechanism operates in macrophages, we stimulated with LPS RAW264.7 macrophages pretreated with siRNA to Sec22b, its binding partner Stx5, or both. In that regard, knockdown of one or both SNAREs resulted in the inhibition of NO release (Figure 5D). Similar to the effect of pharmacological inhibitors on iNOS expression, Sec22b knockdown diminished iNOS protein levels (Figure 5E), as well as *Inos, Tnf* and *Il6* expression, as assessed by quantitative reverse transcription PCR (RT-qPCR) (Figure 5F).

**Figure 5.**
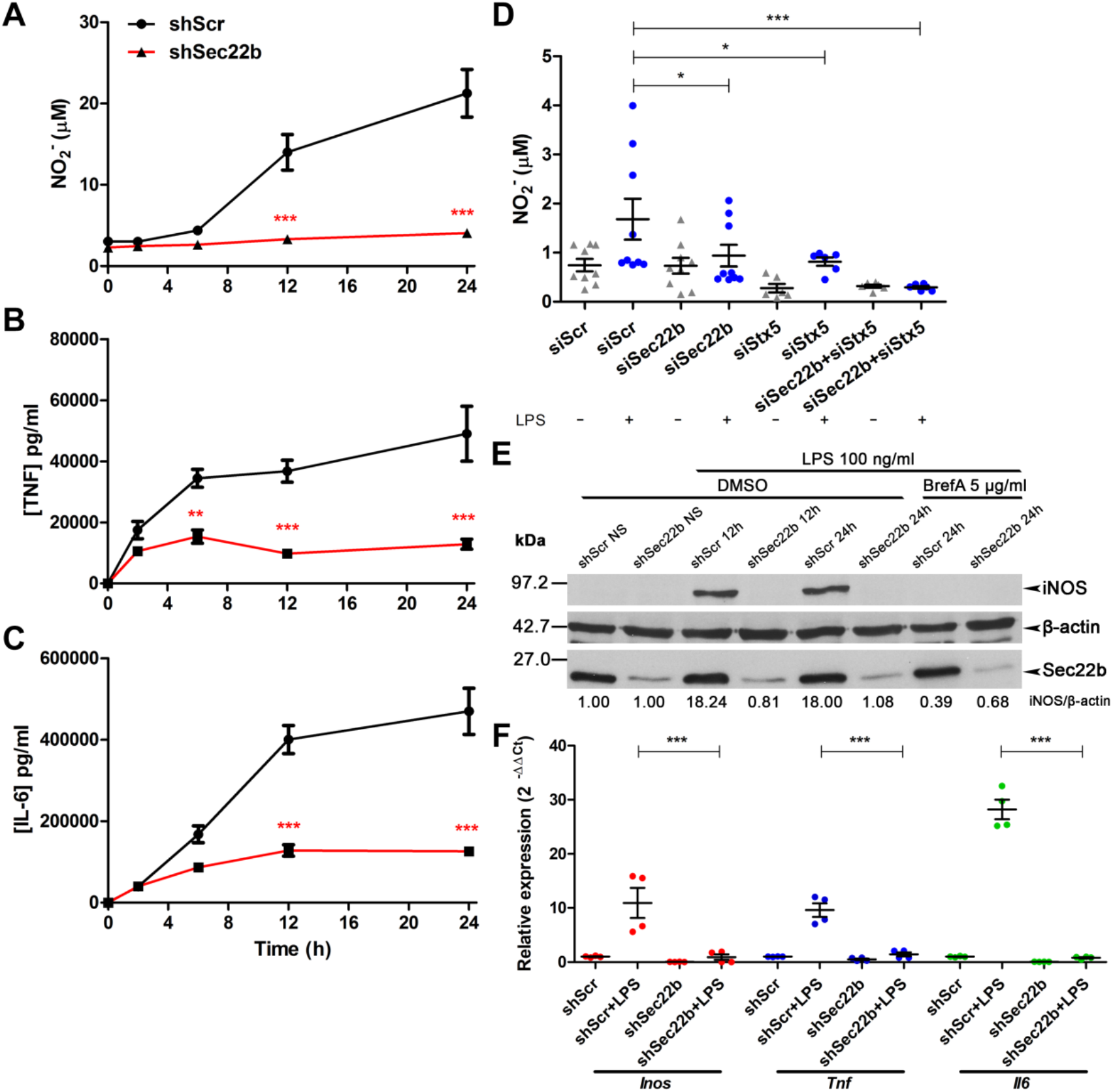
Sec22b is a positive regulator of inflammatory mediator release. To evaluate whether Sec22b modulates inflammatory effector secretion, LPS was used to stimulate JAWS-II cells transduced with an shRNA targeting Sec22b (shSec22b) or a scrambled sequence (shScr). Over a period of 24 h, the release of NO **(A)** TNF **(B)** and IL-6 **(C)** was quantified in cell culture supernatants. Data are presented as mean concentration ± SEM of three independent experiments done in triplicate wells. **(D)** NO secretion in control or LPS-treated RAW264.7 cells transfected with siRNA to a scrambled sequence, Sec22b or Stx5. Data are presented as mean concentration ± SEM of at least two independent experiments done in triplicate wells; each point represents the concentration of a well. **(E)** iNOS protein levels in non-stimulated (NS) or LpS-stimulated Sec22b-KD JAWS-II cells preincubated with DMSO or brefeldin A (BrefA) were assayed via Western blot. iNOS densitometries were normalized with those of the β-actin loading control, and expressed relative to NS-DMSO-treated cells. Images are representative of two experiments. **(F)** *Inos, Tnf* and *Il6* mRNA levels in LPS-stimulated Sec22b-KD JAWS-II cells were assayed via RT-qPCR. Data are presented as mean relative expression ± SEM of two independent experiments done in duplicate wells; points represent relative expression in each well. *, *p* < 0.05; **, *p* < 0.01; ***, *p* < 0.001; ns, not significant.

To ensure that the function of Sec22b on inflammatory mediator production is not contingent on LPS stimulation, we treated Sec22b-KD cells with agonists to TLR2 (peptidoglycan [PGN]), dectin-1/TLR2 (zymosan), TLR2 (*L. major* promastigotes), TLR7 (imiquimod) or TLR9 (CpG). Twenty-four hours post-stimulation, silencing Sec22b abrogated the induction of NO, TNF and IL-6 by all of the tested ligands (Figure 6A-C). Because pharmacological blockade of the secretory pathway abrogated inflammatory mediator release (Figure 5), it is possible that the resident SNARE Sec22b is responsible for this abrogation. To test this, we treated shScr and shSec22b cells with brefeldin A, monensin and retro-2 and assessed inflammatory mediator release post-LPS stimulation. Our data showed that inhibitor treatment and Sec22b knockdown did not have a cumulative effect on NO, TNF and IL-6 secretion (Figure 6D-F), thereby implying that Sec22b is largely responsible for this phenotype.

**Figure 6.**
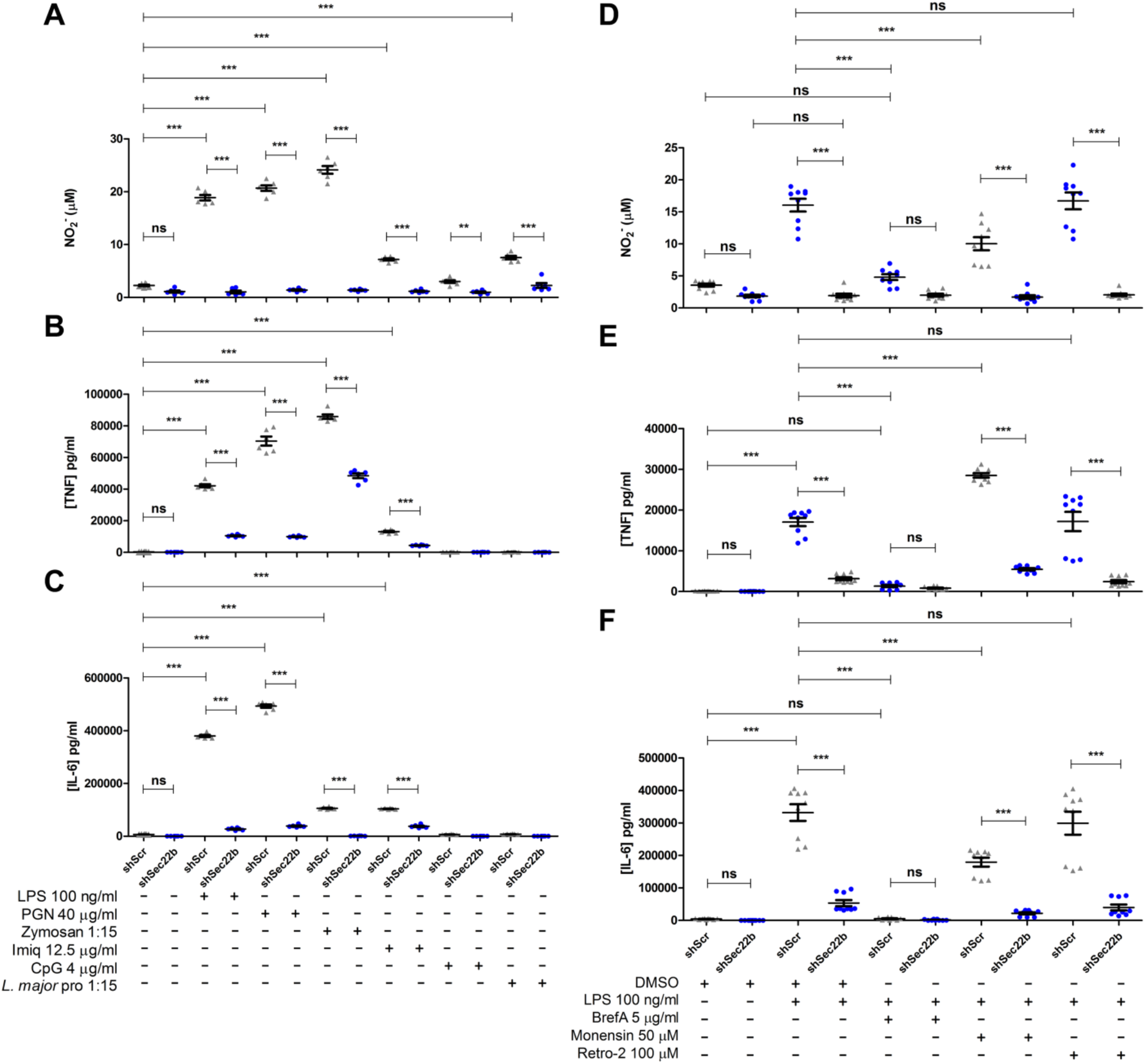
The role of Sec22b is neither specific to LPS stimulation, nor cumulative with chemical disruption of the secretory pathway. **(A)** To evaluate whether Sec22b modulates inflammatory effector secretion in an exclusively LPS-dependent manner, shScr- or shSec22b-transduced JAWS-II cells were stimulated with TLR ligands LPS, PGN, zymosan, imiquimod, CpG and *L. major* promastigotes for 24 h. Cell culture supernatants were then probed for the release of NO **(A)** TNF **(B)** and IL-6 **(C)**. To show whether ER-Golgi disruption had a cumulative effect with Sec22b knockdown, JAWS-II cells were treated with pharmacological inhibitors brefeldin A (BrefA), monensin or retro-2. Cell culture supernatants were then probed for the release of NO **(D)** TNF **(E)** and IL-6 **(F)**. In all panels, data are presented as mean concentration ± SEM of at least two independent experiments done in triplicate wells; each point represents the concentration of a well. **, *p* < 0.01; ***, *p* < 0.001; ns, not significant.

### Sec22b participates in the nuclear translocation of NF-κB

To understand how Sec22b regulates LPS-induced responses, we evaluated the impact of Sec22b knockdown on LPS-induced signal transduction cascades leading to the expression of iNOS. Apart from an impairment of JNK phosphorylation, silencing of Sec22b in JAWS-II DC had no significant effect on LPS-induced signaling events and on NF-κB expression (Figure 7A). To determine whether Sec22b is involved in the nuclear translocation of NF-κB, we stimulated JAWS-II cells expressing either a shScr or a shSec22b with LPS and quantified NF-κB nuclear translocation using imaging flow cytometry (Figure 7B-C) and immunoflurescence confocal microscopy (Figure 7D-E). Our results show that knockdown of Sec22b hampered LPS-induced NF-κB nuclear translocation. We made a similar observation in LPS-stimulated RAW264.7 macrophages treated with siRNA to Sec22b (Figure 7F-G). These data indicate that the secretory pathway-resident SNARE Sec22b modulates proinflammatory effector production by participating to the nuclear translocation of NF-κB.

**Figure 7.**
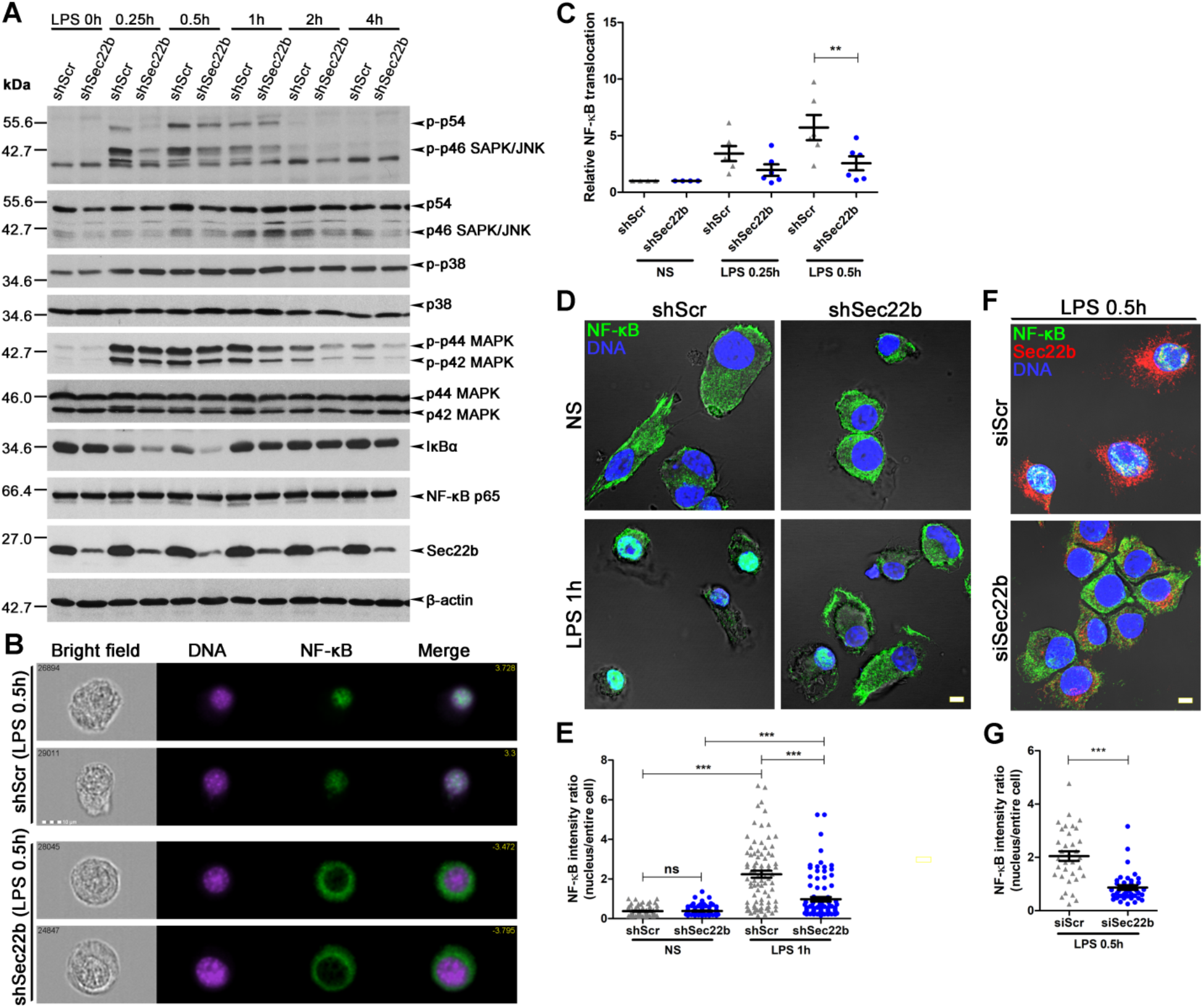
Sec22b modulates MAPK phosphorylation and the translocation of NF-κB to the nucleus. **(A)** ShScr- or shSec22b-transduced JAWS-II cells were stimulated with LPS for 0.25-4 h. Western blots show levels of total and phosphorylated MAPK, as well as IκBα and NF-κB. β-actin was used as loading control and images are representative of two experiments. Nuclear translocation of NF-κB (green) was visualized via imaging flow cytometry **(B)** and quantified **(C)**. DNA is in magenta; bar, 10 μm. Data are presented as average NF-κB translocation ± SEM relative to non-stimulated (NS) cells of at least five independent experiments; each point represents one experiment. NF-κB translocation was also visualized and quantified via confocal microscopy in control and LPS-stimulated JAWS-II cells **(D** and **E)** or RAW264.7 macrophages **(F** and **G)**. DNA is in blue; bar, 5 μm. Data are presented as average MFI ratios (NF-κB MFI of nucleus/entire cell) ± SEM of four independent experiments (≥10 cells per experiment); each point represents the ratio of a single cell. **, *p* < 0.01; ***, *p* < 0.001; ns, not significant.

Since Sec22b is present in the ER-ERGIC-Golgi circuitry (8,29), we hypothesized that NF-κB and Sec22b are found in the same co partments. To test this, we used immunofluorescence to examine the colocalization of NF-κB with Sec22b, as well as other ERGIC and Golgi proteins. In control and LPS-stimulated BMDC, we found partial colocalization under steady state conditions, which disappeared following LPS-induced migration of NF-κB to the nucleus (Figure 8A-B and Figures S2 & S3). We also observed that Sec22b was present in the nucleus; LPS slightly, but significantly, increased the proportion of Sec22b in the nucleus compared to the entire cell (Figure 8C). Similarly, immunogold electron microscopy revealed that NF-κB and Sec22b cooccur in membranous structures in the cytoplasm, and increasingly in the nucleus after LPS stimulation (Figure 8D and Figure S3). Both NF-κB and Sec22b were found in nuclear regions that do not co-stain with DAPI. Since nucleoli also exhibit this characteristic (30), we asked whether Sec22b colocalizes with nucleolar marker fibrillarin (31), and found that Sec22b was not found in these structures (Figure 8E-F and Videos S1 & S2). Collectively, these findings revealed that NF-κB is an ER-Golgi-associated protein that necessitates Sec22b to translocate to the nucleus.

**Figure 8.**
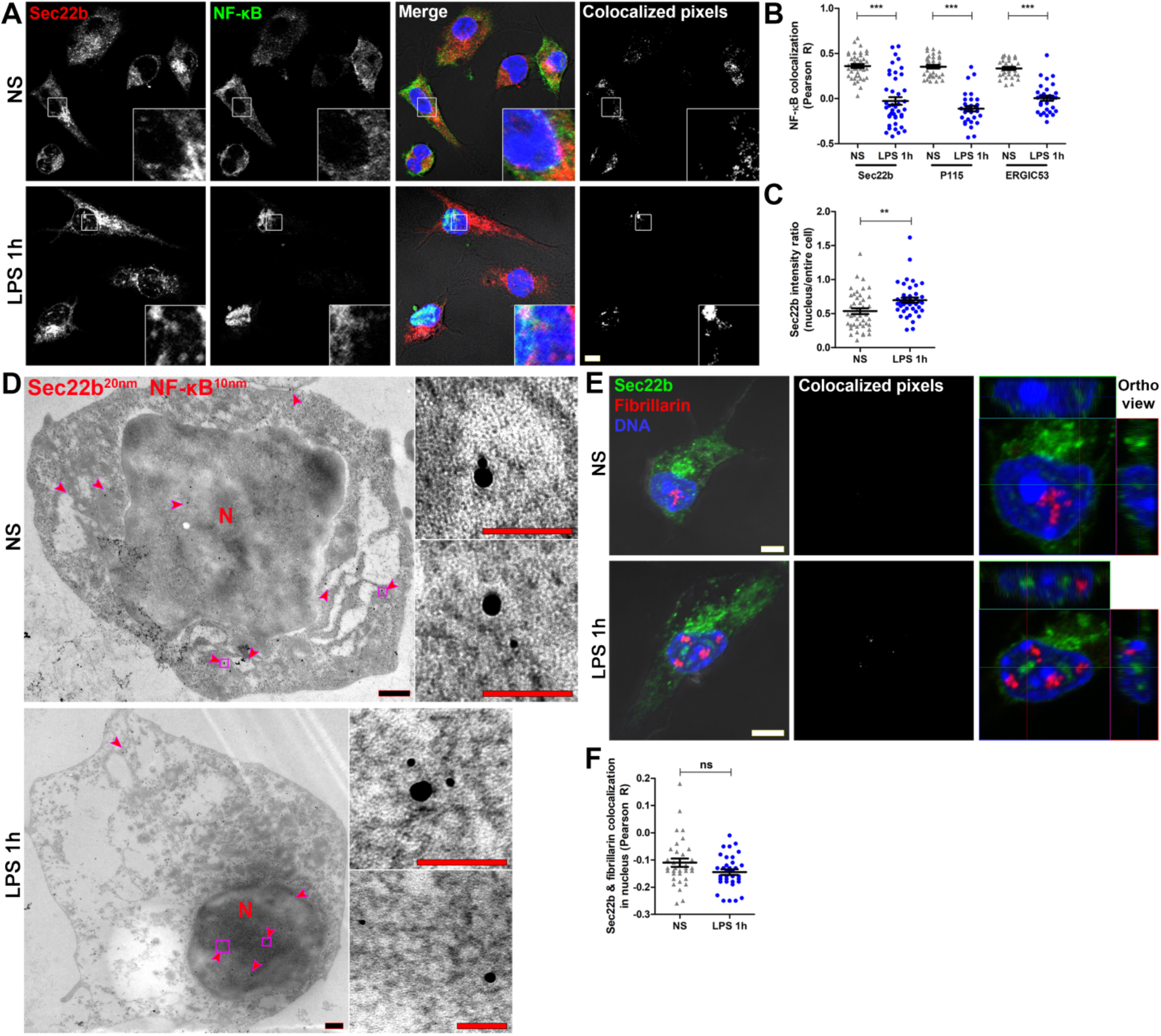
NF-κB and Sec22b co-migrate to the nucleus. **(A)** BMDC were stimulated with LPS for 1 h and the colocalization (white pixels, rightmost panels) of NF-κB (green) with SNARE Sec22b (red) was assessed by immunofluorescence. DNA is in blue; 3.5X-enlarged insets are shown. **(B)** NF-κB colocalization was quantified using the Pearson method (see also Figure S2). **(C)** Quantification of Sec22b MFI ratio in nucleus relative to the entire cell. **(D)** Immuno-electron microscopy assessment of NF-κB (10 nm nanoparticles) and Sec22b (20 nm nanoparticles) expression in non-stimulated (NS) and LPS-treated BMDC. Red arrowheads indicate regions where both NF-κB and Sec22b nanoparticles were at a ≤100 nm distance (see also Figure S3). **(E)** Assessment of Sec22b (green) colocalization with nucleolar marker fibrillarin (red). Rightmost panels show orthogonal views of the nuclear region (see also Videos S1 & S2). **(F)** Quantification Sec22b and fibrillarin colocalization in the nucleus. Image panels are representative of three independent experiments. Scale bars denote 5 μm (white), 500 nm (black) and 100 nm (red). In (B), (C) and (F), data are presented as mean ± SEM of three independent experiments (≥10 cells per experiment); each point represents the measurement of a single cell. **, *p* < 0.01; ***, *p* < 0.001; ns, not significant.

## Discussion

In dendritic cells, microbial-derived molecules induce the expression of genes encoding pro-inflammatory mediators and anti-microbial molecules through complex signalling cascades that culminate in the activation of specific transcription factors (32). In the present study, we aimed at further elucidating the mechanisms and pathways associated to the production of NO and inflammatory mediators in BMDC exposed to LPS. Our main finding is that the secretory pathway, through the action of the SNARE Sec22b, plays a central role in LPS-induced responses by regulating the nuclear translocation of the transcription factor NF-κB.

Whereas the signaling pathways leading to the expression of iNOS and the roles of NO have been explored in great details in various immune cell types (16,18), much less is known on the sub-cellular localization of iNOS. Using electron microscopy and biochemical approaches, Vodovotz and colleagues reported that over 40% of the iNOS activity in whole sonicates was particulate in LPS-activated primary macrophages (33). They also observed that iNOS was associated to 50 to 80 nm vesicles of undefined nature, which did not correspond to endosomes, lysosomes or peroxysomes. Interestingly, the most intense iNOS labeling observed by immuno-electron microscopy was perinuclear, on the trans side of the trans-Golgi network (33). Upon phagocytosis, iNOS was found to associate to phagosomes through an actin-dependent mechanism (34). Our results provide evidence that in dendritic cells stimulated with LPS, iNOS is present within the secretory pathway, as it colocalizes to various extents with proteins found in the ER, the ERGIC and the Golgi. Moreover, we observed that on phagosomes, iNOS colocalized with ER-Golgi SNAREs Sec22b and Stx5, raising the possibility that the previously observed iNOS-positive vesicular structures originate from these organelles.

Given the large number of genes controlled by NF-κB, this ubiquitous transcription factor plays a central role in the regulation of inflammation, and innate and adaptive immune responses, including LPS-induced responses in dendritic cells (32,35). In resting cells, NF-κB dimers are sequestrated in an inactive state in the cytoplasm by a member of the inhibitory proteins known as IκBs (36), preventing NF-κB from being recognized by the nuclear import machinery (37). Cell stimulation triggers signaling cascades leading to the phosphorylation of IκB, its ubiquitination, and degradation by the proteasome (38,39), enabling the nuclear import of NF-κB dimers, their binding to consensus DNA sequences, and the activation of target gene expression (25). Nucleocytoplasmic trafficking of macromolecules such as transcription factors is tightly controlled by importins and exportins, through the recognition of nuclear localization signals and nuclear export signals (40). Using pharmacological inhibitors, we obtained evidence that the ER-Golgi circuitry contributes to optimal LPS-induced responses in BMDC, through the regulation of MAPK phosphorylation and NF-κB nuclear translocation. The fact that Erk1/2 and p38 are involved in LPS-induced NO release (41) led us to the finding that secretory pathway blockade inhibited Erk1/2 and p38 phosphorylation, which may explain in part why NO secretion decreases. Consistent with our findings, previous studies with RAW264.7 macrophages and with primary mouse astrocytes revealed that ER stress induced by either tunicamycin or brefeldin A inhibited LPS-induced iNOS expression (42,43). These findings highlight a previously unappreciated role for the ER-Golgi secretory pathway in the translocation of NF-κB to the nucleus.

Through their role in membrane fusion, the ER/ ERGIC-resident SNAREs Sec22b and Stx5 participate in various cellular processes including secretory autophagy (28), embryonic development (15), and plasma membrane expansion (44). These two SNAREs also play a key role in phagosome maturation and function by regulating the delivery of ER and ERGIC resident proteins to phagosomes in macrophages and dendritic cells. Remarkably, we found that Sec22b was needed for the expression, at the mRNA and protein levels, of iNOS, TNF and IL-6. In immune cells, LPS-induced responses are largely dependent on NF-κB, as it regulates the expression of several genes including iNOS, IL-6, and TNF (25). Although Sec22b KD also decreased MAPK phosphorylation, its effect on NF-κB translocation was surprising but not without precedent. For one, ER- and Golgi-resident proteins such as HSP90 and golgin-97 have been implicated in the translocation of NF-κB to the nucleus (45,46). Since we observed that NF-κB was present in the ERGIC and Golgi, it is reasonable to postulate that Sec22b plays an active role in the cytoplasmic-nuclear shuttling of NF-κB. Second, SNAREs and their chaperones are required for nuclear pore assembly and sealing (47). Our finding that Sec22b and NF-κB co-occured in the cytoplasm and nucleus further supports the notion that Sec22b is part of the circuity that, with help from importins (37,40), shepherds NF-κB translocation.

There is growing evidence that SNAREs and alternatively spliced isoforms access the nucleus where they perform unsuspected functions (47–52). In a study aimed at determining the subcellular localization of Stx17, it was found that this SNARE is present in both the cytoplasm and nucleus of several cell types (52). The absence of homology with known nuclear localization signal (NLS) consensus sequences led the authors to speculate about the existence of a yet uncharacterized nuclear transport mechanism for Stx17. The meaning of the presence of Stx17 in the nucleus remains to be elucidated. Similarly, the SNARE SNAP-47, was found in the ER, ERGIC and nucleus of HeLa cells (50). Further analyses revealed the presence of several putative NLS and two nuclear export signals in SNAP-47. Treatment of HeLa cells with leptomycin B, which inhibits CRM-1/ exportin1-dependent nuclear export led to nuclear accumulation of SNAP-47, suggesting a functional nuclear export signal in SNAP-47 (50). SNAP-47 was previously shown to interact with Stx3 (49). Interestingly, a splice variant of Stx3 was found to undergo nuclear translocation, to bind to several transcription factors, and act as a transcriptional activator (48). An isoform of Stx1, lacking a transmenbrane domain, was also found to localize in the nucleus (51). The observation that Sec22b was present in non-nucleolar compartments of the nucleus raises the interesting possibility that Sec22b partakes in nuclear dynamics and gene regulation. Since the ‘a’, ‘b’ and ‘c’ isoforms of Sec22 possess transmembrane domains (29), we do not exclude the possibility that Sec22b undergoes cleavage prior to its nuclear translocation. Hence, how Sec22b migrates to the nucleus is subject of future research.

In sum, our findings demonstrated that the SNARE Sec22b controls inflammatory effector release by facilitating the translocation of NF-κB from the secretory pathway to the nucleus.

## Experimental procedures

### Ethics statement

Mouse work was done as stipulated by protocol 1706-06, which was approved by the *Comité Institutional de Protection des Animaux* of the INRS-Centre Armand-Frappier Santé Biotechnologie. This protocol respects procedures on animal practice promulgated by the Canadian Council on Animal Care, described in the Guide to the Care and Use of Experimental Animals.

### Antibodies and inhibitors

Rabbit and mouse antibodies anti-iNOS, -SAPK/ JNK, -p-SAPK/JNK (Thr183/Tyr185), -p38, -p- p38 (Thr180/182), -p44/42 MAPK, -p-p44/42 MAPK (Thr202/Tyr204), -IκBα, and -NF-κB were purchased from Cell Signaling [cat. nos. 13120, 2982, 9252, 9251, 9212S, 9211, 9102, 9101, 9242, 4764, 6956]; rabbit antibodies anti-Sec22b and -Stx5 from Synaptic Systems [186003, 110053]; rabbit anti-ERGIC53/p58 and mouse anti-β-actin from Sigma [E1031, A1978]; rabbit anti-PDI and -CNX from Enzo Life Sciences [ADISPA890D, ADISPA860D]; rabbit anti-fibrillarin from Abcam [ab4566]; and rabbit anti-P115 from Proteintech [135091AP]. Pharmacological inhibitors brefeldin A (diluted to 0.5 μg/ml), monensin (5 μM) and retro-2 (100 μM) were obtained from Molecular Probes [B7450], Sigma [M5273] and Calbiochem [554715], respectively, and were reconstituted in DMSO from Bioshop [DMS555].

### Cell culture

Bone marrow-derived dendritic cells (BMDC) were differentiated from the bone marrow of 6-to 8-week old female C57BL/6 mice. BMDC were differentiated for 7 days in RPMI (Life technologies) containing 10% v/v heat-inactivated foetal bovine serum (FBS) [Life Technologies] 10 mM HEPES (Bioshop) at pH 7.4, penicillin-streptomycin (Life Technologies), and 10% v/v X63 cell-conditioned medium as a source of granulocyte-macrophage colony-stimulating factor (53,54). Experiments were carried out in complete medium containing 5% X63 cell-conditioned medium. JAWS-II dendritic cell-like lines stably transduced with shRNA for Sec22b (10) were a gift from D. S. Amigorena (Institut Curie). JAWS-II cells were cultured in RPMI containing 20% heat-inactivated FBS, 10 mM HEPES at pH 7.4, penicillin-streptomycin, 10% X63 cell-conditioned medium and 40 μg/ ml puromycin (Bioshop). The mouse macrophage cell line RAW264.7 clone D3 was cultured in DMEM containing L-glutamine (Life Technologies), 10% heat-inactivated FBS, 10 mM HEPES at pH 7.4, and penicillin-streptomycin. For microscopy experiments, BMDC and JAWS-II cells were seeded onto poly-L-lysine coated coverslips (BD Biosciences, 354085) and RAW264.7 cells onto uncoated glass coverslips (Fisher, 1254581). All mammalian cells were kept in a humidified 37°C incubator with 5% CO_2_.

*Leishmania major* NIH S (MHOM/SN/74/ Seidman) clone A2 (A2WF) promastigotes were provided by W. R. McMaster (University of British Columbia) and were cultured in *Leishmania* medium [M199-1X (Sigma) with 10% heat-inactivated FBS, 40 mM HEPES at pH 7.4, 100 μM hypoxanthine, 5 μM hemin, 3 μM biopterin, 1 μM biotin, and penicillinstreptomycin] in a 26°C incubator. Freshly differentiated promastigotes in late stationary phase (5-day cultures at >50×10^6^ promastigotes/ml) were used for infections.

### siRNA knockdown

RAW264.7 macrophages in the second passage were seeded onto 24-well plates and reverse-transfected with Lipofectamine RNAiMAX (Thermo Fisher Scientific, 13778075) as per the manufacturer’s recommendations. Cells were transfected with siRNAs targeting a scrambled sequence, Sec22b (Thermo Fisher Scientific D0018101005 and 4390815) or Stx5 (Dharmacon, L063346010005) at a final concentration of 25 nM in a 600 μl volume of complete DMEM without antibiotics for 48 h (55). Prior to experiments, macrophages were replenished with complete medium for 4 h before lipopolysaccharide (LPS) [diluted to 100 ng/ml; Sigma, L3129] stimulation. Cells were subsequently prepared for microscopy, and their culture supernatants used for NO measurements.

### Phagocytosis assays

Cells were incubated at 4°C for 5 min with mouse serum-opsonized zymosan (Sigma, Z4250) or *L. major* promastigotes, followed by a 2 min spin at 298.2 *g* in a Sorvall RT7 centrifuge (56). Particle internalization was triggering by transferring cells to 37°C, and after 15 min (zymosan) or 2 h (promastigotes), excess particles were washed thrice with warm medium. Cells were then lysed or prepared for confocal microscopy (56).

### NO and cytokine release measurements

Cells were pretreated for 3 h with DMSO or the aforementioned chemical inhibitors and subsequently stimulated - in the presence or absence of inhibitors - with LPS, peptidoglycan (PGN) [40 μg/ml; Sigma, 77140], Imiquimod [12.5 μg/ml; Invivogen, tlrl-imq], CpG [4 μg/ml; Invivogen, tlrl-1585], zymosan or *L. major* promastigotes. Cell culture supernatants were spun to remove debris, and nitrite concentrations were assayed via the Griess reaction (57). ELISA kits were used to quantify secreted TNF (Ready-SET-Go! Mouse TNFα Kit, eBiosciences,88732488) and IL-6 (BD OptiEIA, Becton-Dickinson Biosciences, 555240), as per the manufacturers’ instructions.

### Immunofluorescence confocal microscopy and imaging flow cytometry

Cells on coverslips were fixed with 2% paraformaldehyde (Thermo Scientific, 28906) for 20 min and blocked and permeabilized for 17 min with a solution of 0.1% Triton X-100, 1% BSA, 6% non-fat milk, 2% goat serum, and 50% FBS. This was followed by a 2 h incubation with primary antibodies diluted in PBS. Then, cells were incubated for 35 min in a solution containing suitable combinations of AlexaFluor-linked secondary antibodies (diluted 1:500; Molecular Probes A11008, A11011, A11031, A11001, A21247) and DAPI (1:40000; Molecular Probes, D3571). G_M1_^+^ lipid rafts were stained with the cholera toxin-subunit B-AlexaFluor 647 conjugate (1:5000; Molecular Probes, C34778) (58). Coverslips were washed three times with PBS after every step. Coverslips were mounted onto glass slides (Fisher) with Fluoromount-G (Southern Biotechnology Associates, 010001) and sealed with nail hardener (Sally Hansen, 45077). Cells were imaged with the 63X objective of an LSM780 confocal microscope (Carl Zeiss Microimaging). Control stainings revealed no cross-reactivity or background (Figure S3). Images were taken in sequential scanning mode via the ZEN 2012 software and mounted with Photoshop (Adobe). Images were analyzed using Icy Software (Institut Pasteur) (59). Colocalization (60) was evaluated using the Pearson R coefficient calculated from regions of interested defined by active contours (61) on the cytoplasm (DIC channel) or on the nucleus (DAPI). Nuclear translocation was evaluated by dividing the mean fluorescence intensity of an entire cell by that of its nucleus. Recruitment was evaluated by scoring the presence of a given protein on the phagosome membrane (58,62).

Imaging flow cytometry experiments were performed as previously described (63). Briefly, cells were washed with cold PBS containing 1% horse serum (Sigma), 0.1% NaN3, and 5 mM EDTA, fixed, permeabilized and stained. DNA was stained with DAPI immediately prior to running the samples through an ImageStreamX MKII flow cytometer running IDEAS software (Amnis). Nuclear translocation of NF-κB was evaluated using a score that quantifies the correlation of NF-κB and DAPI pixels on a per cell basis; a score of >1 indicated nuclear translocation (63).

### Immunogold labeling and electron microscopy

LPS-treated treated or control BMDCs were fixed overnight at 4°C in 0.1% glutaraldehyde + 4% paraformaldehyde in a cocodylate buffer at pH 7.2. After washing, cells were treated with 1.3% osmium tetroxide in collidine buffer and dehydrated (56). Pelleted cells were embedded (SPURR, TedPella) and placed in BEEM capsules (Pelco Int, 130). After resin polymerization, samples were sectioned using an ultramicrotome (LKB Broom - 2128, Ultratome). Sections were placed on nickel grids, treated with sodium metaperiodate and blocked with 1% BSA in PBS1X. Grids were then incubated with primary antibodies for 90 min (diluted 1:30 with 0.1% BSA in PBS1X), washed, and incubated in suitable 10 nm (anti-mouse) or 20 nm (anti-rabbit) gold particle-conjugated secondary antibodies (Abcam ab39619 and ab27237) for 60 min (diluted 1:10 with 0.1% BSA in PBS1X). Washed samples were treated with uranyl acetate and lead citrate for contrast, and imaged via a Hitachi 7100 electron microscope mounted with an AMT XR-111 camera. Control stainings revealed no background or non-specificity (Figure S3).

### Electrophoresis and Western blotting

Cells were washed with cold PBS containing 1 mM Na_3_VO_4_ (Sigma) and lysed in a solution containing 1% NP-40, 50 mM Tris-HCl (pH 7.5), 150 mM NaCl, 1 mM EDTA (pH 8), 1.5 mM EGTA, 1 mM Na_3_VO_4_, 50 mM NaF, 10 mM Na_4_P_2_O_7_, and complete protease inhibitors (Roche) (56). After incubation at −70°C, lysates were centrifuged for 15 min to remove insoluble material. After protein quantification, 30 μg of sample was boiled (100°C) for 6 min in SDS sample buffer and migrated in SDS-PAGE gels. Proteins were transferred onto Hybond-ECL membranes (Amersham Biosciences, 10600003), blocked for 2 h in TBS1X-0.1% Tween containing 5% BSA, incubated with primary antibodies (diluted in TBS1X-0.1% Tween containing 5% BSA) overnight at 4°C, and thence with suitable HRP-conjugated secondary antibodies (GE Healthcare NA931V and NA934V) for 1 h at room temperature. Washed membranes were incubated in ECL (GE Healthcare, RPN2106) and immunodetection was achieved via chemiluminescence (62). Densitometric analysis of Western blot bands was done using the AlphaEase FC software (Alpha Innotech) with β-actin used as loading control.

### RNA extractions and RT-qPCR

RNA from BMDC and JAWS-II cells was extracted using TRIzol (Life Technologies, 15596026) as per the manufacturer’s protocol. Purified RNA was reverse-transcribed (64) and cDNA was submitted to RT-qPCR analysis using the iTaq Universal SYBR Green Supermix (BioRad, 1725121). Samples were amplified in duplicate on a MX3000P instrument (Stratagene), and the following primers were used. *Inos:* 5’-CTGCTGGTGGTGACAAGCACATTT-3’ (AD529F) and 5’-ATGTCATGAGCAAAGGCGCAGAAC-3’ (AD530R); *Tnf:* 5’-GACGTGGAAGTGGCAGAAGAG-3’ (AD537F) and 5’-TGCCACAAGCAGGAATGAGA-3’ (AD538R); *Il6:* 5’-ACAACCACGGCCTTCCCTACTT-3’ (AD539F) and 5’-CACGATTTCCCAGAGAACATGTG-3’ (AD540R); and *Hprt:* 5 ‘ - GTTGGATACAGGCCAGACTTTGTTG-3’ (AD55F) and 5’-GATTCAACTTGCGCTCATCTTAGGC-3’ (AD56R). MxPro-MX2000P software (Stratagene) was used to record cycle threshold (C_*T*_) values, which were analyzed by the 2^-ΔΔC*T*^ method. Data were normalized to *Hprt* and expressed in terms of fold change differences relative to non-stimulated cells.

## Data analysis

Statistical differences between two or multiple groups were assessed via the Mann-Whitney *U* test or one/two-way ANOVA followed by Bonferroni post-hoc tests, respectively. Two-way ANOVA was used to evaluate time-dependent differences in inflammatory mediator release. Differences were considered significant when *p* < 0.05 and graphs were constructed using GraphPad Prism 5.0 (GraphPad Software Inc).

## Data availability

All data are contained within the article and its supporting information.

## Acknowledgements

We are grateful to D. S. Amigorena for the kind gift of shRNA-transduced JAWS-II cells, W. R. McMaster for providing *L. major* promastigotes, J. Tremblay for assistance in immunofluorescence experiments, and A. Nakamura for assistance in electron microscopy.

## Funding and additional information

This work was supported by Natural Science and Engineering Research Council of Canada grant 1055043 to AD and by Canadian Institutes of Health Research (CIHR) grant PJT-159647 to SS. AD is the holder of the Canada Research Chair on the Biology of intracellular parasitism. GAD was partially supported by a CIHR Frederick Banting and Charles Best Doctoral Award and by bridging funds from the Centre for Host-Parasite Interactions.

## Author contributions

Conceived and designed the experiments: GAD, RD, and AD. Performed the experiments: GAD, RD, JD and AF. Analyzed the data: GAD, RD, JD, AF, SS and AD. Contributed reagents/materials/analysis tools: SS. Wrote the paper: GAD and AD. All authors discussed the findings and commented on the manuscript.

## Conflict of Interest

The authors declare no conflicts of interest in regards to this manuscript.

